# Noncytotoxic bacterial nano-melanin with immense cosmetic potential produced by *Bacillus infantis* isolated from salt pans of Marakkanam, Tamil Nadu, India

**DOI:** 10.64898/2026.09.17.752272

**Authors:** S Mohamed Ansar, Noble K Kurian

**Affiliations:** School of Life Sciences, BS Abdur Rahman Institute of Science and Technology, Vandalur, Chennai; Department of Research & Development, TIBR Biotech Private Limited, Kalamassery Kochi, 682022, India

**Keywords:** Melanin, bacteria, SPF, cosmetic, anti-ageing

## Abstract

Melanin is a high molecular weight pigment derived from the oxidation of phenols, followed by the polymerization of phenols and the quinones they produce. Usually, they appear in dark brown or black color. *Bacillus infantis* was isolated from saltpans of Marakkanam, Tamilnadu. This strain was utilized for melanin production by screening using tyrosine basal broth, and the extracted pigment was characterized by UV-visible spectrometer and FT-IR. Notably, melanin was found to be nano size and it is confirmed by DLS analysis. The nano-sized melanin-producing strain *Bacillus infantis* was confirmed as non-pathogenic by testing with different antibiotics. This study also explores the sun protection factor of melanin, and this melanin shows successful outcomes. The antioxidant activity of melanin shows a wide range of radical scavenging activity by DPPH, ABTS and FRAP assays. The melanin has shown modest anti-ageing activity at the higher concentration tested. Nano melanin was found to be non-cytotoxic even in higher concentrations tested. This is the first report on melanin production by *Bacillus infantis*, and there is limited information available to date about melanin-producing bacteria from salt pan sediments.

## Introduction

Melanins are ubiquitous phenolic pigments found in most of the living organisms. The pigment is responsible for the survival of life on Earth as it is widely accepted that it has an exceptional capacity to absorb a broad spectrum of radiation. (Tran-Ly et al.,2020). The pigment is high in molecular weight and formed with various structural properties produced through oxidative polymerization with quinones. It is ubiquitous in nature, and even very old fossils of dinosaurs, early birds, non-avian theropod taxa, and primitive cephalopods have been proven to contain melanin. (Kurian et al., 2015).

The classes of melanin include eumelanin, pheomelanin, pheomelanin, neuromelanin, and allomelanin, which are determined by the polymerization pathways, building blocks, and enzymes involved. Depending on the class, Melanin has different properties. Dihydroxyindole and DHICA combine to form the most prevalent form of eumelanin or DOPA (o-dihydroxyphenylalanine) and is in the color of brown to black and produced by a variety of microbes, including fungus and bacteria. UV radiation triggers melanogenesis in human skin, which can result in the formation of skin melanin and the yellowish pheomelanin that makes up human skin. Furthermore, several catechol moieties in fruits and plants have been linked to the synthesis of allomelanin or DHN (di-hydroxynaphthalene). The brain chemical neuromelanin is essential for the treatment of neurodegenerative diseases (Choi et al., 2021).

It should be noted that synthetic melanin cannot be compared to natural melanin. Natural melanin is not only safer but also more cost-effective than its artificial counterpart. However, microorganisms have recently been explored as an alternative to chemical melanin production. Microorganisms offer a more reliable method for producing melanins compared to Sepia and vegetable melanin. Unlike those methods, microorganisms are not affected by seasonal fluctuations during production and can modify their mechanisms according to the composition of the medium and growth conditions provided (El-Naggar et al., 2022).

Microbial melanin has an important advantage in that it can be derived from various bacterial and fungal sources, each with unique characteristics and applications. In laboratory conditions, researchers have identified and optimized microbial species that have the capacity to produce melanin (El-Naggar et al., 2022). Several types of bacteria that have been identified from the natural world, including *Aeromonas, Bacillus, Rhizobium, and Streptomyces*, have been shown to use tyrosinase to make melanin. Several other bacteria like *Pseudomonas stutzeri, Pseudoalteromonas lipolytica* etc. shown to produce pyomelanin produced via homogentisic acid. *Cellulophaga tyrosinoxydans* produces a yellow pigment that is suggested to be a pheomelanin (Kurian, 2015). Properties of melanin vary slightly among the melanins while general physicochemical characteristics remain common.

Melanin pigment can now be useful in many fields, including material science, biomedicine, environmental remediation, and cosmetics. From a physiochemical perspective, Melanin functions as a broad UV visible light spectrum absorber in nature. This pigment not only blocks UV light but also has strong antioxidant properties. Melanin also demonstrates semiconductor-like activity that is reliant on hydration. It is, therefore, assessed by a part of organic electronic devices. The bioavailability, biocompatibility, and biodegradability of microbial melanin are further benefits that make it a viable option for biomedical application and radiation protection applications (Tran-Ly et al.,2020).

Several studies have reported melanin-producing bacteria from marine sources and salt desert regions (Gordhanbhai et al., 2022; Tarangini et al., 2013). This indicates that salt stress should be one of the factors triggering melanin biosynthesis pathways. Melanin-producing bacteria from salt pan sediments have been less explored so far. In this study, we mainly focus on melanin-producing bacteria from the saltpan regions of Tamil Nadu, India.

## Materials and Methods

### Sample collection

The soil sample was collected from Marakkanam(12.2024°N, 79.9567°E), Tamil Nadu, India. Soil samples were collected from the wetland of a salt pan in sterile polyethylene bags, which we brought into the laboratory on ice. The samples were kept briefly in refrigerated condition before it is plated into nutrient agar plates.

### Isolation of bacteria from the soil sample

The soil samples were serially diluted and plated on nutrient agar plates with 4% sodium chloride. The plates were incubated at 37oC for 48 hours. Colonies having unique morphology and pigmentation were selected for purification. Purification was done using quadrant streaking of the colonies in nutrient agar plates with 4% NaCl and incubating for 24 hours. Pure colonies from the last quadrant are picked and stored temporarily in a nutrient agar medium with 4% NaCl at 4°C.

### Screening for melanin producers

The isolated pure colonies were screened for melanin production in tyrosine basal broth (Yabuuchi & Ohyama, 1972) with 2g/L L-tyrosine as the sole source of carbon and nitrogen. 2% of isolates and basal broth were transferred to 100mL medium, and 0.2% of L-tyrosine was added separately to the broth after autoclaving. The production flasks were incubated for 8-10 days at 37oC. The color change in the medium was monitored daily.

### Characterization of melanin-producing strain CM03

#### Microscopy and biochemical characterization

The strain CM03 is gram-stained to identify its gram nature. IMViC and catalase tests were performed as biochemical tests to support the identification of bacteria up to the species level. (Faddin et al., 1976).

### Antibiotic Susceptibility test and MAR index

The Kirby-Bauer disc diffusion method was used to profile antibiotic sensitivity (Bauer et al., 1966). On a Mueller-Hinton agar (HiMedia, India) plate, a consistent bacterial smear was created using a sterile cotton swab. Each plate could only contain four antibiotic discs. The discs were placed on the plate. Space was created between the discs to allow the inhibition zone’s growth. Prior to the examination, the plates were incubated for twenty-four hours at 37°C. Based on the manufacturer’s provided (Himedia, India) information about the size of the inhibition zones surrounding each disc, the outcome was classified as resistant, intermediate, or sensitive. (Performance standard for antimicrobial disc susceptibility tests, 2006). The formula a/b, where “a” is the number of antibiotics to which the isolate was resistant and “b” is the number of antibiotics to which the isolate was exposed, was used to determine the MAR index (Krumperman.,1983).

### Evaluation of Halotolerance of CM03 strain

Bacteria classified as halotolerant can thrive in high salt concentration environments and conditions where salt is not present. For 24 to 48 hours at 37°C, the halotolerance test of the Melanin producing strain CM03 was streaked on nutritional agar plates containing 0.5%, 2%, 4%, 6%, 8%, 10%, and 15% NaCl (Grigary et al., 2024). Plates were incubated for up to 48 hours, and the growth of the bacteria in the plates was observed.

### Molecular characterization of strain CM03

Isolates were identified at the National Centre for Microbial Resource (NCMR) DNA sequencing facility, National Centre for Cell Science, Pune. At the facility, genomic DNA was isolated by the standard phenol/chloroform extraction method, followed by PCR amplification of the 16S rRNA gene using universal primers 16F27 [5’-CCA GAG TTT GAT CMT GGC TCA G-3’] and 16R1492 [5’-TAC GGY TAC CTT GTT ACG ACT T-3’]. The amplified 16S rRNA gene PCR product was purified by PEG-NaCl precipitation and directly sequenced on an ABI® 3730XL automated DNA sequencer (Applied Biosystems, Inc., Foster City, CA) as per manufacturer’s instructions. The sequencing was performed from both ends using additional internal primers to read each position at least twice. Assembly was carried out using Lasergene package followed by identification using the EzBioCloud database

### Phylogenetic Analysis of Strain CM03

The nucleotide sequences were all converted to FASTA format, and the Clustal W algorithm (Thompson et al., 1997) in BioEdit software was used to perform multiple sequence alignment for the assembled nucleotide sequences (Hall, 1999). For additional analysis, aligned sequences were loaded into a MEGA5: Molecular Evolutionary Genetics Analysis (MEGA) software version 5.0 (Tamura et al., 2007). The alignment’s ends were cut to ensure that each sequence had the same length, and the aligned sequences were then put into MEGA format in order to do phylogenetic analysis. Utilizing the nucleotide-based TN84 evolutionary model for the construction of the phylogenetic tree, the neighbor-joining approach (Saitou & Nei, 1987) is used to estimate genetic distances based on synonymous and nonsynonymous nucleotide changes. We estimated the statistical support for branching using 1000 bootstrap steps.

### Extraction and Purification of melanin

The cell-free supernatant from the tyrosine basal broth was acidified to pH 2 using 1 N HCl after the mixture was incubated for around 48 hours. At the bottom of the flask, when the pH drops, a black precipitate of melanin is visible. After allowing the precipitate to rest for 2 days at room temperature (RT) to precipitate fully, it was centrifuged. The final black pellet was rinsed 2 times each using 15 mL of 0.1 N HCl, ethanol, and then water, respectively. The mixture was then utilized for further analysis after being lyophilized for 6 hours (Sajjan et al., 2013). The purity of melanin is tested by thin-layer chromatography. The extracted pigment was spotted on a TLC plate and subjected to a solvent system including n-butanol, acetic acid, and water (12:3:5). The spots were developed by spraying ninhydrin reagent and kept in a hot air oven for 3-5 minutes.

### Characterization of the purified pigment

#### Chemical solubility test

The pigment was confirmed as melanin by the soluble nature of the melanin, tested with selected solvents: distilled water, Sodium hydroxide, ethanol, Isopropanol, Methanol, Acetone, N-butanol, Dimethyl sulfoxide, Benzene, Phenol, and Chloroform. Solvents were added to melanin vortexed for 5 min and observed for solubility.

### UV visible spectrophotometry

A UV-visible spectrum was produced by scanning a 100 µg/mL melanin solution diluted in 0.1 N NaOH at 200 and 550 nm wavelengths. In the analysis, 0.1 N of NaOH was used as the blank. To verify the pigment’s authenticity, the spectrum produced was compared to a standard of 100 µg/mL synthetic melanin (Yuan et al., 2007).

### FT-IR Spectrophotometry

The purified bacteria melanin was pressed into discs at high pressure using a pellet maker and IR grade KBr (1:10). A Thermo Nicolet Avatar 370 spectrophotometer with a KBr beam splitter and a DTGS (Deuterated Triglycine Sulphate) detector (7800-350 cm-1) at the Instrumentation Center of the BS Abdur Rahman Institute of Science and Technology, Vandalur, Chennai, Tamilnadu, was used to record the FT-IR spectrum at 4,000-400 cm-1 and resolution 4 cm-1 (Ravishankar et al., 1995).

### Dynamic Light Scattering (DLS) Analysis and Zeta Potential

The sample was measured after dispersion in Milli-Q water and sonication for 10 min before the measurement. Dispersed melanin samples were observed in the concentration of 100µL/1mL. Size distribution and surface charge of a melanin powder were determined by the dynamic light scattering (DLS) and zeta potential instrument (Zetasizer Nano ZS, Malvern) at 25 °C

### Cosmetic potential of the pigment

#### Sun Protection Factor (SPF) Estimation

The capacity of melanin to raise the Sun Protection Factor (SPF). 1 mL of 100% ethanol with 100µg/mL concentration of melanin powder is used. Using ethanol as the blank, the mixture’s absorbance in the UV range (290–320 nm) was measured at intervals of 5 nm. SPFs were calculated using the following formula by Mansur et al. (1986), where CF (correction factor) = 10; EE (l) = arrhythmogenic effect of radiation with wavelength k; Abs (l) = spectrophotometric absorbance value of the solution; and I = solar intensity spectrum. EE (l) ×I is constant and was determined.

### Antioxidant activity of melanin

#### DPPH radical scavenging assay

Ascorbic acid was used as the reference standard for the DPPH assay. The ascorbic acid stock solution was prepared in distilled water (1 mg/ ml; w/v). A 60μM solution of DPPH in methanol was freshly prepared, and a 200µl of this solution was mixed with 50μl of the CM03 melanin at various concentrations (1.56, 3.12, 6.25, 12.5, 25, 50,100,200,400,800 and 1000 µg/ml). The plates were kept in the dark for 15 minutes at room temperature, and the absorbance decreased at 515 nm. Control was prepared with DPPH solution only, without melanin or ascorbic acid. 95% methanol was used as blank.

The following formula calculated radical scavenging activity

Antioxidant activity (%) = Absorbance of Control-Absorbance of test/ Absorbance of control ×100

### ABTS antioxidant assay

The reaction was initiated by the addition of 200µl of diluted ABTS to 1.56-1000 µg/ml of different concentrations of melanin, and in control, 50 µl of methanol was used instead of the sample. Methanol is used as a blank. The absorbance was read at 734 nm, and the percentage of antioxidant activity was calculated. The inhibition was calculated according to the equation:

% of antioxidant activity =A0-A1/A0×100

Where A0 is the absorbance of the control, and A1 is the absorbance of the test compound.

### Ferric Reducing Antioxidant Power (FRAP) Assay

The FRAP assay was performed as described by Nishaa et al. (2012), with minor modifications. The FRAP working reagent was freshly prepared by mixing 2.5 mL of 10 mM 2,4,6-tris(2-pyridyl)-s-triazine (TPTZ) in 40 mM HCl with 2.5 mL of 20 mM FeCl₃ in 25 mL of 0.1 M acetate buffer (pH 3.6) and incubated at 37 °C for 10 min before use. Sample CMO3 melanin was tested at final concentrations of 6.25, 12.5, 25, 50 was mixed with 2 mL of FRAP reagent. Following 30 min incubation, absorbance was measured at 593 nm against the blank. All measurements were performed in triplicate.

### Collagenase Enzyme Inhibitory (Anti-Ageing) Activity

#### Cell culture and treatment

Human skin melanoma (SK-MEL) cells were seeded in 6-well plates at a density of 0.3 × 10⁶ cells/well and allowed to acclimatize for 24 h under standard culture conditions (37 °C, 5% CO₂). The test sample (CMO3 melanin) and the reference standard, epigallocatechin gallate (EGCG), were prepared in Dulbecco’s Modified Eagle Medium (DMEM) at a stock concentration of 100 mg/mL and sterilized by filtration through a 0.2 µm Millipore syringe filter. Working solutions were then serially diluted in DMEM and added to the cultured cells to give final concentrations of 6.25, 12.5, 25, 50 and 100 µg/mL. Untreated wells served as the negative control. All treatments were performed in biological triplicate, and plates were incubated for a further 24 h before processing.

### Cell lysate preparation

Following treatment, culture media were aspirated and adherent cells were gently washed with pre-cooled phosphate-buffered saline (PBS). Cells were lysed with RIPA buffer and the lysate was clarified by centrifugation at 1000 × g for 5 min. The resulting supernatant was used immediately for the collagenase inhibition assay or stored in aliquots at ≤ −20 °C until analysis.

### Collagenase inhibition assay

Collagenase activity in the cell lysate was measured using a Human Collagenase ELISA kit (Origin, Cat. No. OPK9586) with a TMB substrate, following the manufacturer’s protocol with minor modifications, based on the methods of Park et al. (2010) and Shanura Fernando et al. (2018). Absorbance was recorded at 450 nm. Percentage inhibition of collagenase activity was calculated as:

Inhibition (%) = [(Absorbancecontrol − Absorbancesample) / Absorbancecontrol] × 100

EGCG, a well-characterized collagenase inhibitor, was run in parallel as the positive control to validate assay performance.

### Determination of IC₅₀

The half-maximal inhibitory concentration (IC₅₀) was derived from the concentration– inhibition relationship. Because a valid IC₅₀ can only be interpolated when inhibition exceeds 50% within the tested concentration range, IC₅₀ was calculable for EGCG but not for sample CMO3, which did not reach 50% inhibition at the highest concentration tested (100 µg/mL).

### Elastase Enzyme Inhibitory (Anti-Ageing) Activity

#### Cell culture and treatment

Human skin melanoma (SK-MEL) cells were seeded in 6-well plates at a density of 0.3 × 10⁶ cells/well and allowed to acclimatize for 24 h under standard culture conditions (37 °C, 5% CO₂). The test sample (CMO3 melanin) and the reference standard, epigallocatechin gallate (EGCG), were prepared in Dulbecco’s Modified Eagle Medium (DMEM) at a stock concentration of 100 mg/mL and sterilized by filtration through a 0.2 µm Millipore syringe filter. Working solutions were serially diluted in DMEM and added to the cultured cells to give final concentrations of 6.25, 12.5, 25, 50 and 100 µg/mL. Untreated wells served as the negative control. All treatments were performed in biological triplicate, and plates were incubated for a further 24 h before processing.

### Cell lysate preparation

Following treatment, culture media were aspirated and adherent cells were gently washed with pre-cooled phosphate-buffered saline (PBS). Cells were lysed with RIPA buffer and the lysate was clarified by centrifugation at 1000 × g for 5 min. The resulting supernatant was used immediately for the elastase inhibition assay or stored in aliquots at ≤ −20 °C until analysis.

### Elastase inhibition assay

Elastase activity was measured using a chromogenic assay based on the methods of Senior et al. (1982) and Sallenave et al. (1998), with minor modifications. Briefly, a Tris-HCl buffer (179 mM, pH 8) was prepared, and the substrate N-succinyl-Ala-Pro-Phe p-nitroanilide was dissolved in this buffer at 1.65 mM. Porcine pancreatic elastase was prepared at 0.34 U/mL in the same buffer. In microplate wells, 20 µL of cell lysate was combined with 100 µL of buffer and 50 µL of substrate, and the mixture was pre-incubated for 5 min at 25 °C. The reaction was initiated by adding 50 µL of enzyme solution, and the liberated p-nitroanilide was measured spectrophotometrically at 405 nm. One unit of elastase activity was defined as the amount of enzyme liberating 1 µmol of p-nitroanilide per minute. Percentage inhibition of elastase activity was calculated as:

Elastase inhibition (%) = (1 − B / A) × 100 where A is enzyme activity in the absence of inhibitor (control) and B is enzyme activity in the presence of inhibitor (sample or EGCG). EGCG, a well-characterized elastase inhibitor, was run in parallel as the positive control to validate assay performance.

### Determination of IC₅₀

The half-maximal inhibitory concentration (IC₅₀) was derived from the concentration– inhibition relationship. Because a valid IC₅₀ can only be interpolated when inhibition exceeds 50% within the tested concentration range, IC₅₀ was calculable for EGCG but not for sample CMO3, which did not reach 50% inhibition at the highest concentration tested (100 µg/mL).

### *in vitro* Cytotoxicity of CM03 melanin

#### Cell lines and maintenance

The L929 – Mouse fibroblast cell line was procured from the National Centre for Cell Sciences (NCCS), Pune, India. The cells were cultured in Dulbecco’s Modified Eagles Medium (DMEM-Himedia), supplemented with 10% heat-inactivated Fetal Bovine Serum (FBS) and 1% antibiotic cocktail containing Penicillin (100 U/ml), Streptomycin (100 μg/ml), and Amphotericin B (2.5 μg/ml). The cells containing TC flasks (25 cm2) were incubated at 37°C at a 5% CO2 environment with humidity in a cell culture incubator (Galaxy® 170, Eppendorf, Germany).

### MTT Assay

The cells (2500 cells/well) were seeded on 96 well plates and allowed to acclimatize to the culture conditions, such as 37°C and 5% CO2 environment in the incubator for 24 hours. The test samples were prepared in DMEM media (10 mg/mL) and filtered and sterilized using a 0.2 μm Millipore syringe filter. The melanin samples were further diluted in DMEM media and added to the wells containing cultured cells at final concentrations of 6.25, 12.5, 25, 50, and 100μg/mL, respectively. Untreated wells were kept in control. After treatment with the test samples, the plates were further incubated for 24 h. After the incubation period, the media from the wells were aspirated and discarded. 100 μL of 0.5 mg/mL MTT solution in PBS was added to the wells. The plates were further incubated for 2 hours for the development of formazan crystals. The supernatant was removed, and 100μL DMSO (100%) was added per well. The absorbance at 570 nm was measured with a microplate reader. Three wells per plate without cells served as blank. All the experiments were done in triplicates.

The cell viability was expressed using the following formula

Percentage of cell viability = Absorbance of treated/Absorbance of control X 100

### Statistical analysis

The Statistical analysis and the graphs were plotted using GraphPad Prism (Ver. 6) and Microsoft Excel computer programs.

## Result and Discussion

### Screening for melanin producers

CM03 strain turned the color of tyrosine basal broth turned black on the 8th day, which was selected for the production of melanin., and the strain was inoculated in a flask containing tyrosine basal broth; the strain started producing melanin from the 5th day, and on the 9^th^ day, complete black color melanin was produced. The production media is white (Figure 1a) in color during inoculation, which turned black in color on the 9^th^ day (Figure 1b)This production pattern was similar to that of the earlier reports (Kurian & Bhat, 2018).

**Figure 1:**
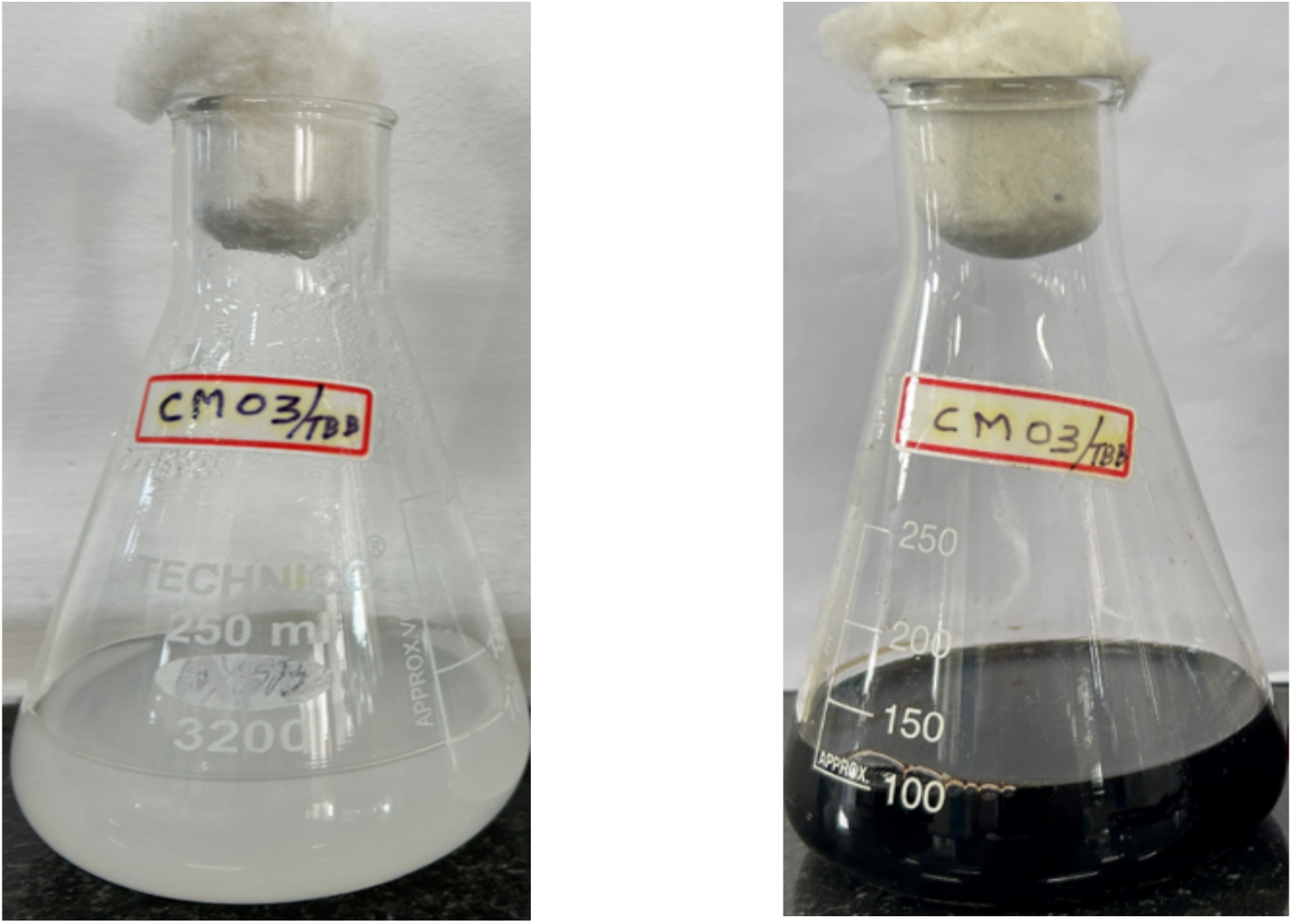
Melanin production by strain CM03 in tyrosine basal broth. (a) media before inoculation (b) media after 9 days of melanin production

### Characterization of melanin-producing strain CM03

#### Gram staining and Biochemical Tests

Through gram staining, the producing CM03 strain was found to be a gram-positive, rod-shaped bacteria. Biochemical characterization shows that strain CM03 can produce indole from tryptophan, which shows indole as positive. The bacterial strain has been found to produce acids by glucose fermentation, showing methyl red as positive. Vogues-Proskauer was found to be negative, while the bacteria utilized citrate in the medium. The strain CM03 was shown to produce a considerable amount of catalase enzymes.

### Antibiotic Susceptibility test and MAR index

Antibiotic susceptibility test was carried out by disc diffusion method, CM03 strain was swabbed over the Muller Hinton agar plate, 11 different antibiotic discs were placed in 3 plates the antibiotic disc forms clear zone against CM03 stain without any contamination, MAR (Multiple Antibiotic Resistance) index was calculated using the formula a/b, where ‘a’ is the number of antibiotics to which the isolate was resistant, and ‘b’ is the number of antibiotics to which the isolate was exposed (Krumperman, 1983). Here, a total of 11 antibiotics were tested, and 1 antibiotic was found to be resistant (Table 1). So, the MAR index is 1/11= 0.09. MAR index values greater than 0.2 indicate high-risk sources of contamination where antibiotics are often used. Our sampling source was confirmed to be antibiotic contamination-free.

**Table 1:** Different antibiotic and their zone of inhibition against CM03 strain.

|  | Resistant | Intermediate | Sensitive |  |  |
| --- | --- | --- | --- | --- | --- |
| Antibiotics | Zone of Inhibition in mm |  |  | CM03 |  |
| Cefoxitin CX 30 mcg | 14 | 15 | 16 | 29 | Sensitive |
| Co-Trimoxazole COT25 25 mcg | 10 | 11 | 16 | 28 | Sensitive |
| Meropenem MRP 10 mcg | 19 | 20 | 23 | 38 | Sensitive |
| Ertapenem ETP 10 mcg | 18 | 21 | 22 | 11 | Resistant |
| Colistin sulfate CS 10 µg |  |  | 11-17 | 15 | Sensitive |
| Ciprofloxacin CIP 5 mcg | 15 | 20 | 21 | 28 | Sensitive |
| Cefuroxime (CXM) 30 mcg | 14 | 17 | 18 | 30 | Sensitive |
| Tigecycline TGC 15 mcg |  |  | 20-27 | 22 | Sensitive |
| Ampicillin/Sulbactam A/S 10/10 mcg | 11 | 14 | 15 | 42 | Sensitive |
| Cefoperazone/ Sulbactam CFS 50/50 mcg |  |  | 28-36 | 37 | Sensitive |
| Ceftizoxime (CZX) 30 mcg | 21 | 22 | 25 | 31 | Sensitive |

### Evaluation of Halotolerance of CM03 strain

A halotolerance investigation was conducted on the chosen isolate CM03. NaCl was added to nutrient agar plates in different concentrations, ranging from 0.5% to 15%.. After 24-48 hours of incubation at 36° C, the growth of the CM03 strain was noted up to a 10% concentration of NaCl (Figure 2). There was no sign of bacterial growth in the 15% NaCl plate. High salt tolerance is one factor that can induce melanin production in microbes (Elsayis et al., 2022). Melanin-producing bacteria were found to be isolated from marine sediments (Kurian & Bhat, 2018) and the salt desert of Kutch Gujarat (Jigna et al., 2022). This halotolerant CM03 could also be utilized to produce melanin pigment.

**Figure 2:**
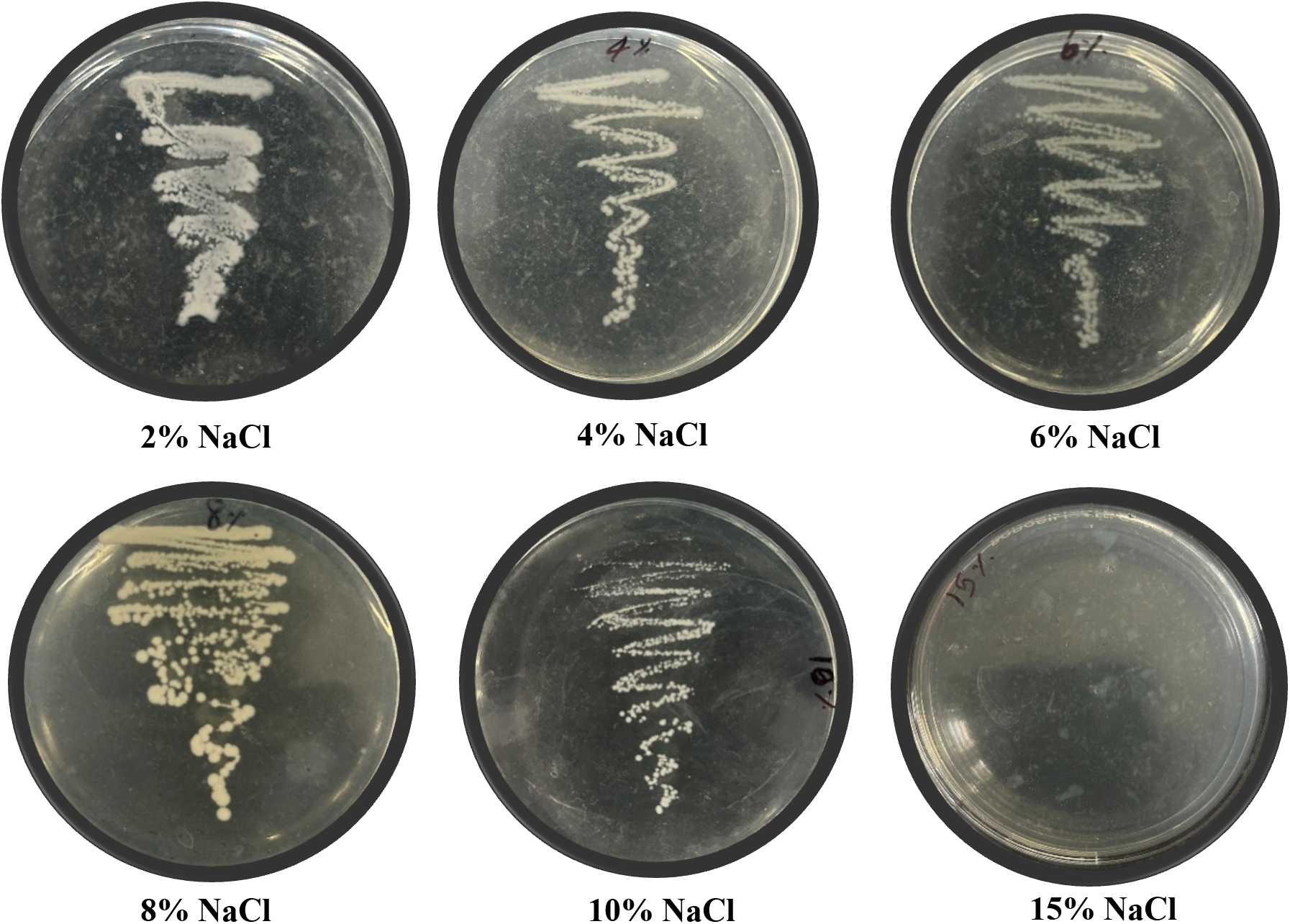
Halotolerance of CM03 strain. Different concentrations of NaCl fed into the Nutrient Agar plate; as the percentage of NaCl increases in the plate, the growth of the CM03 strain decreases gradually

### Molecular characterization of strain CM03

The 16SrDNA sequence is searched against the GenBank database using the Nucleotide BLAST tool. The strain had shown 99.55% similarity with *Bacillus infantis* NNRL B-14911, which confirmed the strain CM03 to be *Bacillus infantis*. The sequence was submitted to NCBI, and an accession number was obtained (CP006643)

### 3.5. Phylogenetic Analysis of Strain CM03

The phylogenetic tree was constructed using 16S rDNA genes from 9 bacterial strains, of which *Pseudomonas aeruginosa* 57 served as the outgroup. *Bacillus infantis* CM03 shared the same node with *Bacillus infantis* strain KKPHNGU1, *Cytobacillus* sp. Strain IS141 and *Neobacillus dentensis* strain CD3. These similar bacteria were found to be halotolerant in nature. The strain had shown less similarity with other Bacillus species like *Bacillus licheniformis, Bacillus subtilis*, etc., as evident from a phylogenetic tree in Figure 3.

**Figure 3:**
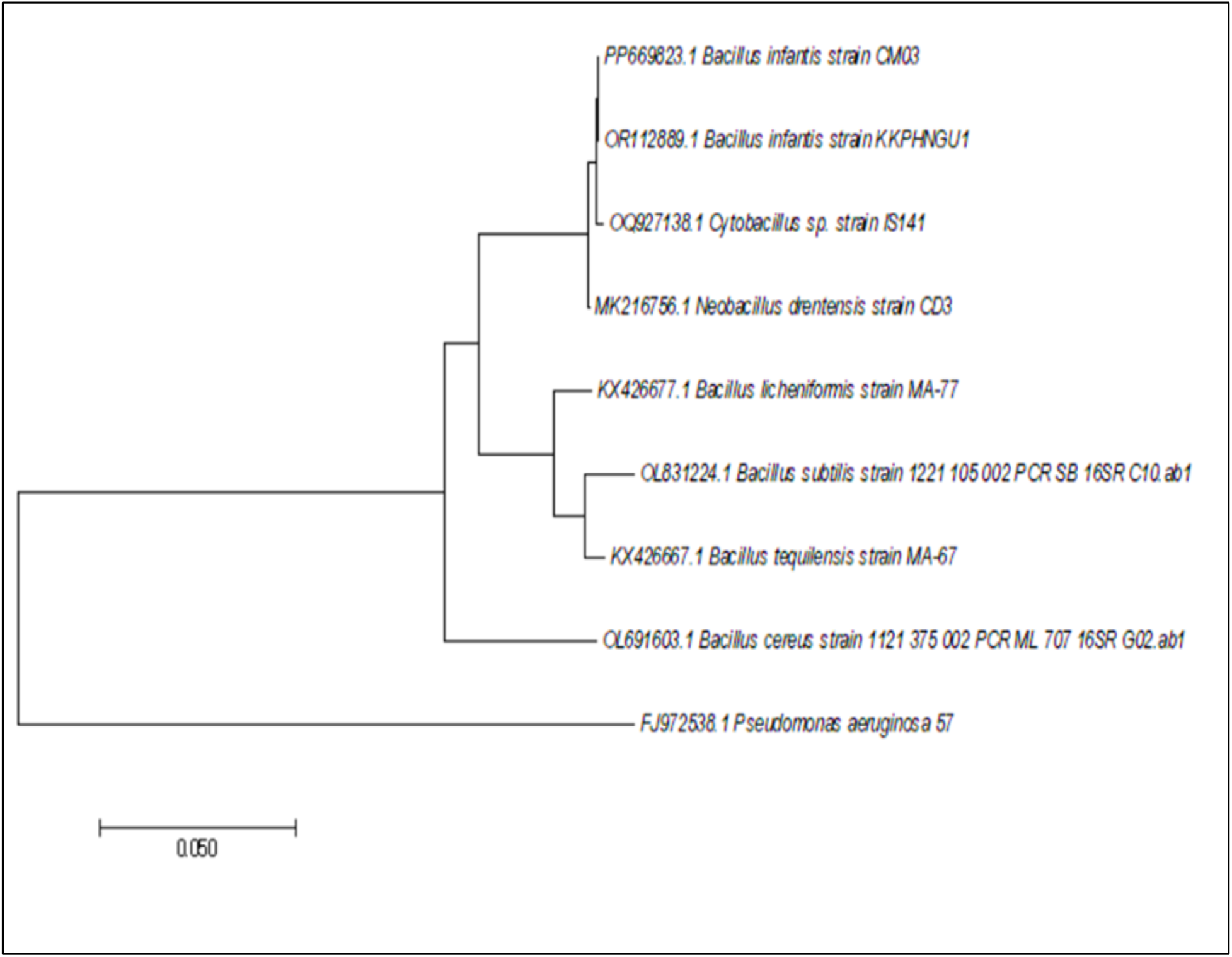
Phylogenetic relationship of *Bacillus infantis* CM03 against other *Bacillus* species.

### Extraction and purification of the pigment

After adjusting the pH of the medium to 2, the precipitate of Melanin pigment was settled down and was observed visibly. Multiple washes with solvents have been done to get fine melanin powder. The purity of the melanin powder was confirmed by getting a single spot without any amino acid contamination using Thin Layer Chromatography (Figure 4). The Rf value was found to be 0.893. The TLC result was found to be similar to earlier reports (Noman et al., 2022; Madhusudhan et al., 2014)

**Figure 4:**
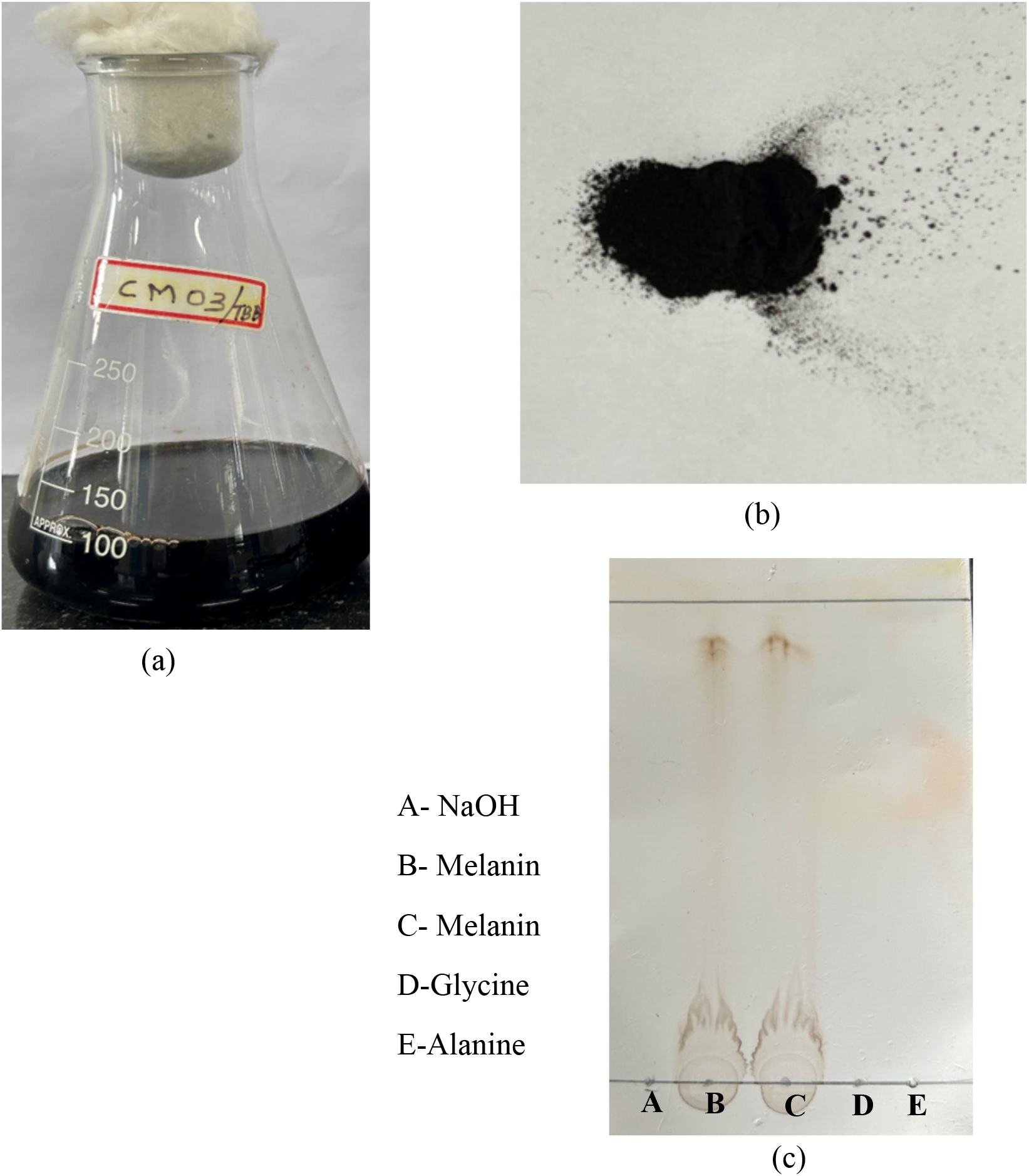
Melanin Extraction and Purification. (a) Medium after the acidification (b)The extracted fine melanin powder after lyophilization and (c) purity confirmation of melanin by thin layer chromatography

### Characterization of the pigment

#### Chemical solubility test

Naturally, melanin is insoluble in water; the pigment was tried to dissolve in water and other chemical solvents. CM03 melanin was found to be completely soluble in sodium hydroxide and Dimethyl sulfoxide. The pigment was found to be partially soluble in methanol, phenol, and ethanol. At acidic pH, the pigment remained insoluble and was found to be insoluble in chloroform, acetone, N-butanol, isopropanol, and benzene. The insolubility in water is used to confirm the pigment as melanin, and the solubility features were found to align with earlier reports (Noman et al., 2022).

### UV visible spectrophotometry

CM03 produced melanin pigment that showed a featureless absorption without an absorption maximum in a UV–visible spectrum analysis. The absorption is higher at 250 nm and decreases gradually at 550 nm (Figure 5). One sharp peak centered approximately at 250 nm, and according to many earlier reports, there are no distinctive absorption peaks in melanin to differentiate it from other cutaneous chromophores. Instead, melanin absorption gradually decreases as wavelength increases from UV to visible spectrum (Mbonyiryivuze et al., 2015)

**Figure 5:**
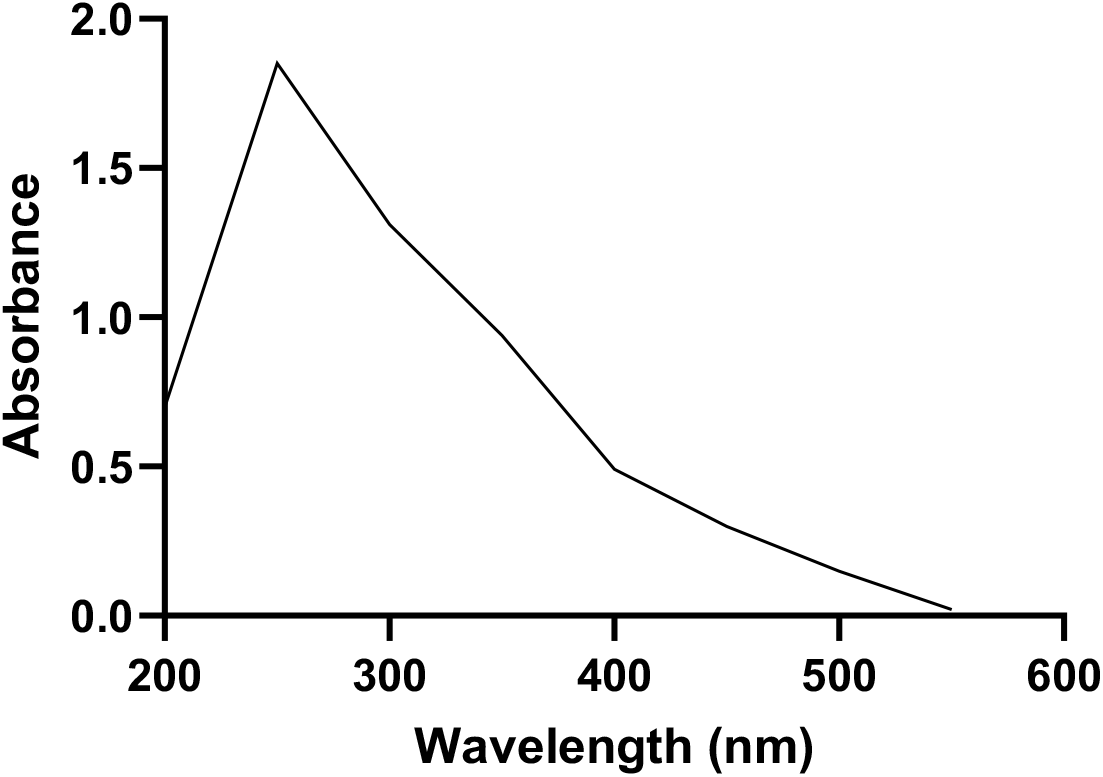
UV-visible spectrum of melanin pigment produced by CM03 strain.

### FT-IR Spectrophotometry

FT-IR spectra of melanin show a broad absorption spectrum at 3266 cm^−1^, which may be due to characteristic O-H stretching or N-H stretching vibrations of the carboxylic acid and phenolic groups in melanin. Characteristic peaks observed in 1606 cm^−1^ and 1436 cm^−1^ were attributed to aromatic ring C = C stretching. The FTIR spectrum showed three spectroscopic curves were fundamentally identical (Figure 6). The spectrum is similar to the earlier reports (Sun et al., 2015).

**Figure 6:**
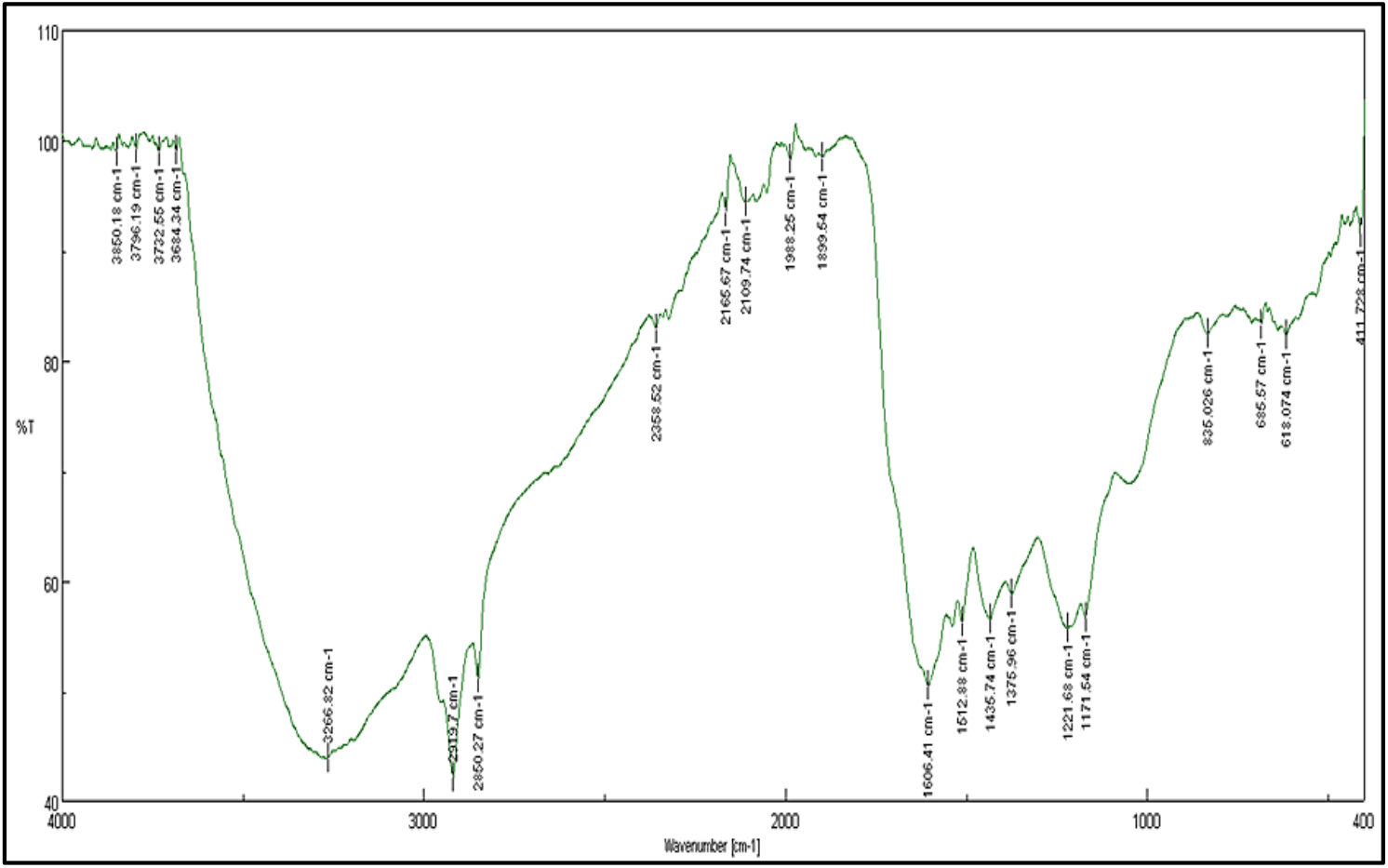
FT-IR Spectrophotometry of melanin pigment produced by CM03 strain.

**Figure 7:**
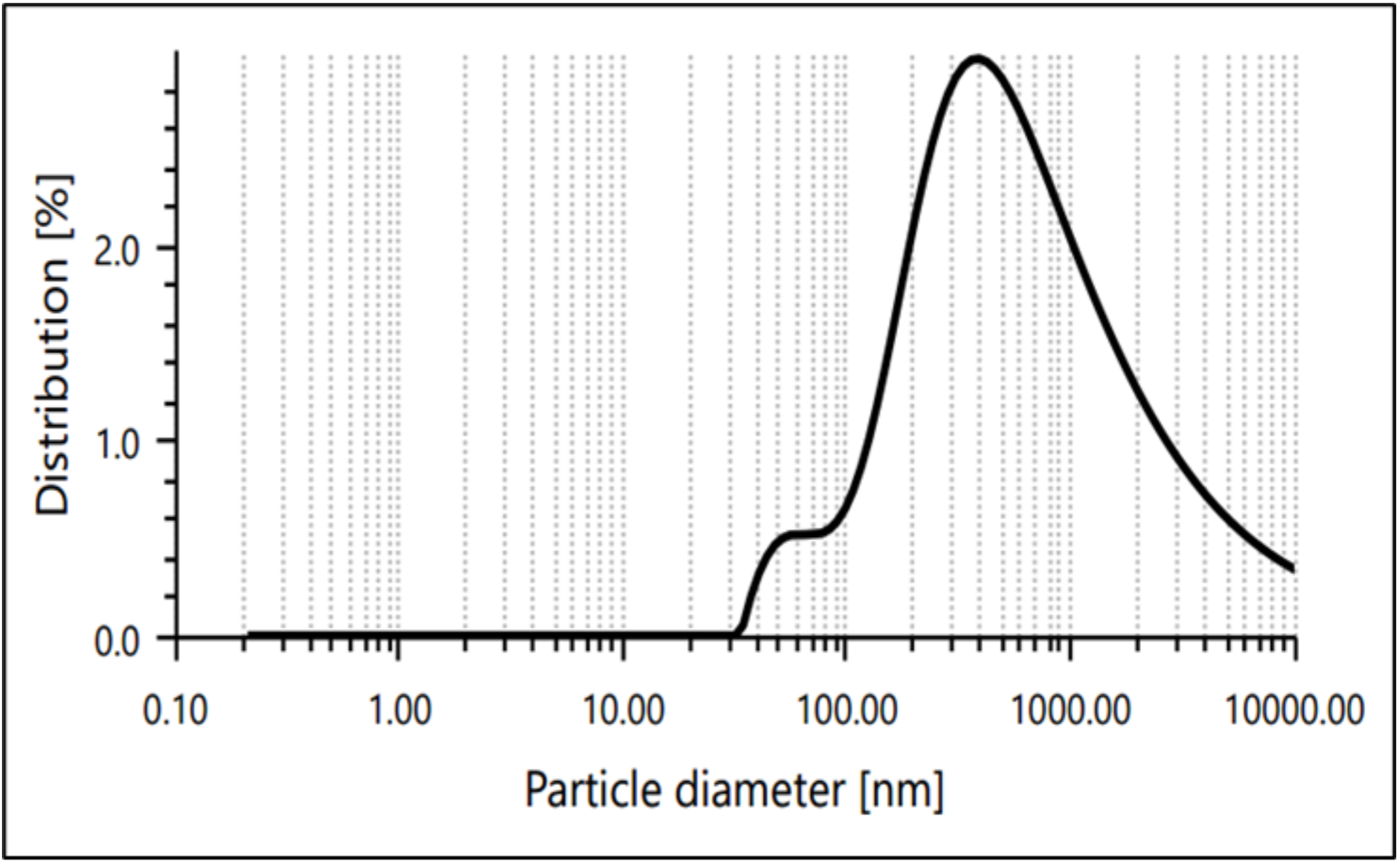
The particle size distribution profile of CM03 melanin.

**Figure 8:**
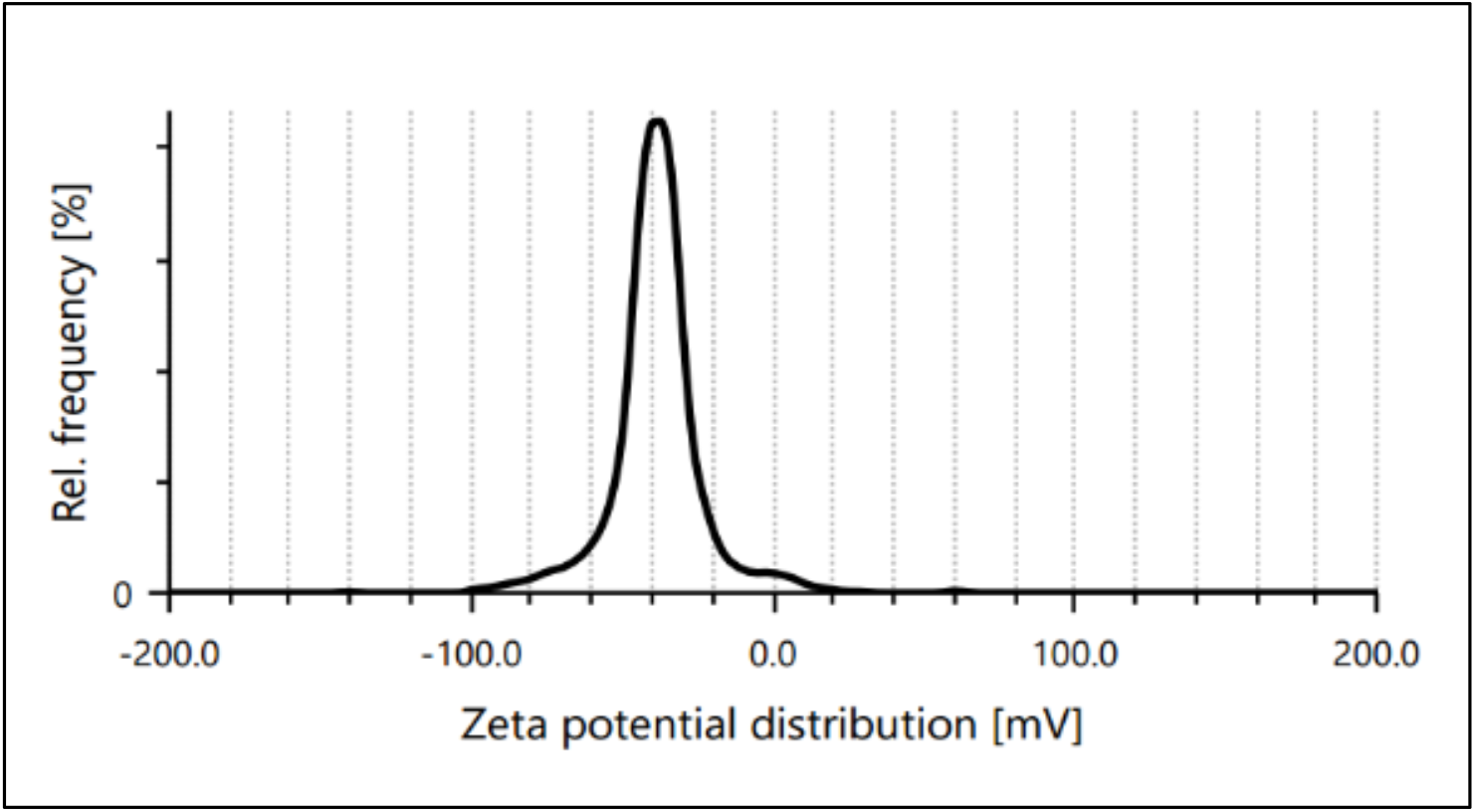
Zeta potential distribution of the melanin nanoparticles.

### Dynamic Light Scattering (DLS) Analysis and Zeta Potential

#### DLS-Particle size analysis

The particle size of melanin particles dispersed in distilled water (by Sonication) was determined using Dynamic Light Scattering analysis. The mean particle size of melanin particles produced by the CM03 strain was 393.9 nm (Figure 12). The CM03 melanin particles were distributed between 52.21 nm and 1173.0 nm. An earlier report in *Pseudomonas sp.* WH00155 showed a similar distribution of melanin nanoparticles with a mean diameter of 519.6nm (Kiran et al., 2017).

### Zeta potential

Zeta potential is an electric potential that is created by the presence of a charge on the surface of a particle. Depending on the chemistry of the particles, zeta potential can be either positive or negative in polarity. It is an indicator of the degree of repulsion between similarly charged particles in a formulation. Repulsive forces prevent particle aggregation during storage, thus making zeta potential indicative of the physical stability of a formulation (Sangwan & Seth, 2022). The zeta potential of melanin particles dispersed in distilled water was found to be −41.2 mV (Figure 13), conferring the stability of CM03 melanin nanoparticles.

### Cosmetic potential of the pigment

#### Sun Protection Factor (SPF) estimation

Sunscreen efficacy is typically measured by the sun protection factor (SPF). This is the ratio of the UV energy required to produce a minimal erythema dose (MED) on protected skin to the UV energy required to produce a MED on unprotected skin. MED refers to the smallest amount of UV light exposure that can cause a perceptible sunburn on unprotected skin. The higher the SPF value, the more protection a sunscreen provides against sunburn (Ebrahimzadeh et al., 2014).

Transmission spectroscopy was utilized to evaluate the *in vitro* activity of the nano-melanin powder. The melanin’s transmission spectrum was measured using a UV-visible spectrometer between 290 and 320 nm, as shown in Table 4. The SPF value of 100µg/mL CM03 nano-melanin was found to be 14.82459±2.51 (Table 2). The distribution of SPF values is classified as follows: SPF 2-4 is minimal, SPF 4-6 is moderate, SPF is 6-8 extra, SPF is maximum 8-15, and SPF>15 is ultra. The higher the concentration, the better the SPF value obtained (Diniatik & Nurulita, 2022). So, The SPF value of nano CM03 melanin was found to be in the ultra-range and could be used in cosmetic formulations.

**Table 2:** The cosmetic potential of the melanin was tested by UV-visible spectroscopy.

| Wavelength | EE(λ)×I(λ) | Trail 1 |  | Trail 2 |  | Mean<br>Abs×EE(λ)×I(λ) |
| --- | --- | --- | --- | --- | --- | --- |
|  |  | Abs | Abs×EE(λ)×I(λ) | Abs | Abs×EE(λ)×I(λ) |  |
| 290 | 0.015 | 1.74 | 0.0261 | 1.37 | 0.02055 | 0.023325±0.003 |
| 295 | 0.0817 | 1.71 | 0.139707 | 1.35 | 0.110295 | 0.125001±0.020 |
| 300 | 0.2874 | 1.69 | 0.485706 | 1.33 | 0.382242 | 0.433974±0.073 |
| 305 | 0.3278 | 1.66 | 0.544148 | 1.31 | 0.429418 | 0.486783±0.081 |
| 310 | 0.1864 | 1.63 | 0.303832 | 1.27 | 0.236728 | 0.27028±0.047 |
| 315 | 0.0837 | 1.59 | 0.133083 | 1.24 | 0.103788 | 0.118436±0.020 |
| 320 | 0.018 | 1.54 | 0.02772 | 1.20 | 0.0216 | 0.02466±0.004 |
|  | **Total** |  | 1.660296 |  | 1.304621 | 1.482459±0.251 |
|  | **SPF** |  | 16.60296 |  | 13.04621 | 14.82459±2.51 |

### Antioxidant activity of melanin nanoparticles

#### DPPH radical scavenging assay

Antioxidants play a critical role in preventing damage caused by free radicals in food and biological systems. Free radicals can accelerate lipid oxidation in food and damage biological macromolecules. The DPPH assay is a reliable method to evaluate the radical scavenging ability of antioxidants. During the test, the antioxidants interact with DPPH radicals and neutralize their free radical character by transferring an electron or hydrogen atom. Figure 15 shows that melanin from the CM03 bacterial strain has shown a significant ability to remove DPPH. With a concentration of 800μg/mL, melanin exhibited 50.11% of radical scavenging activity with an IC50 value of 735.82μg/mL (Figure 9). The control Ascorbic acid used was shown to have 94.95% antioxidant activity, and there were significant differences between melanin and ascorbic acid in the concentrations tested (p < 0.05).

**Figure 9:**
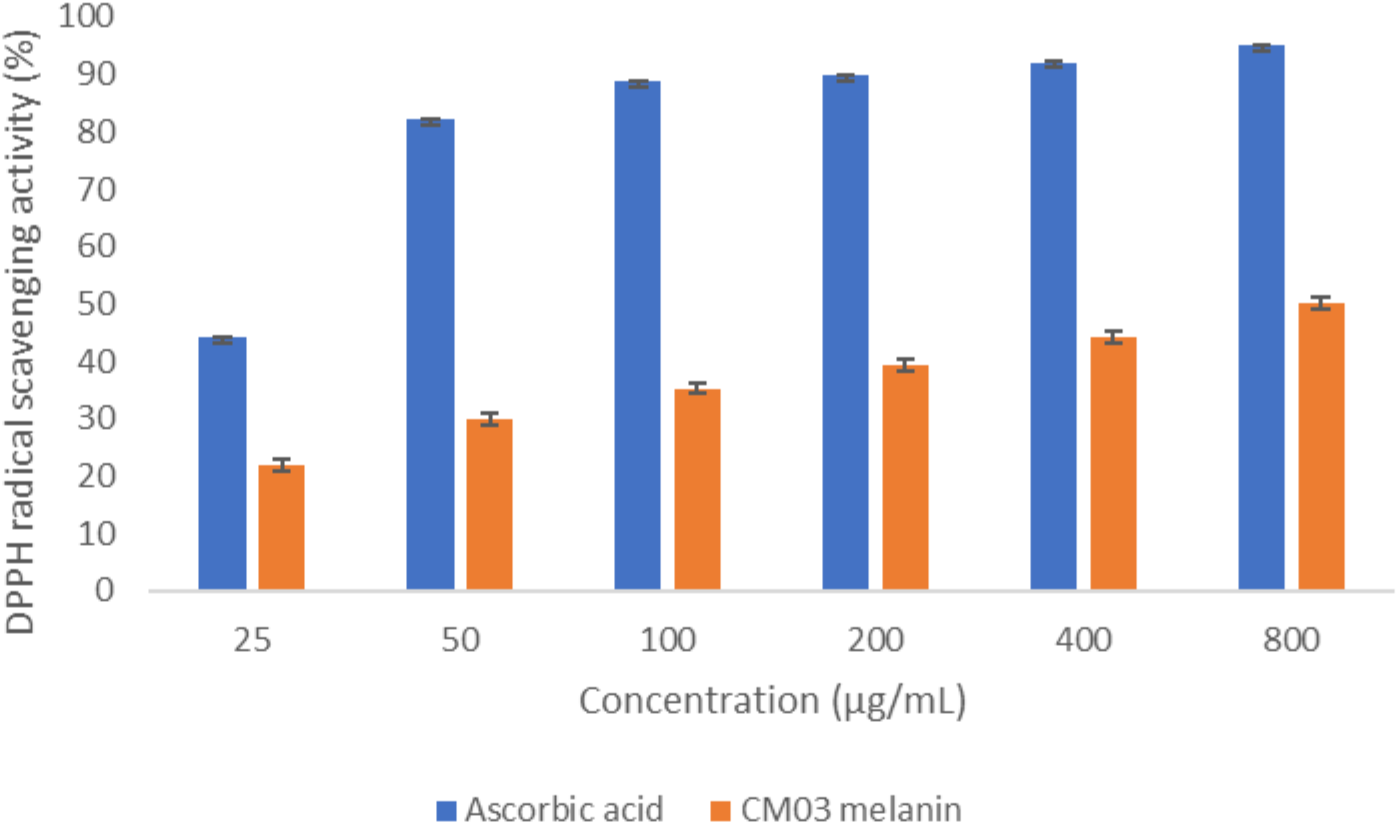
DPPH radical scavenging activity of CM03 melanin.

### ABTS antioxidant assay

The ability of the CM03 melanin to scavenge free radicals was evaluated by inhibiting the oxidation of 2, 2’-azino-bis (3-ethylbenzothiazoline-6-sulfonic acid) (ABTS). The results showed that the purified melanin pigment of strain CM03 showed good antioxidant activity.

The results revealed that 1000μg/mL melanin exhibited a percentage inhibition of 60.73% radical scavenging activity with an IC_50_ value of 447.50μg/mL (Figure 10), comparable to standard antioxidant ascorbic acid, showing activity of 97.23% at the same concentration. There were significant differences between melanin and ascorbic acid in the concentrations tested (p < 0.05). Interaction of melanin pigment with ABTS+ transfers hydrogen atoms to ABTS+, thus neutralizing its free radical character. Due to the unpaired electrons in its molecules, melanin pigment readily interacts with free radicals and other reactive species. It acts as an antioxidant, suggesting its use as a raw cosmetic material to minimize toxin-induced tissue destruction. Melanin interacts with free radicals via simple one-electron transfer processes (El-Naggar & El-Ewasy., 2017).

**Figure 10:**
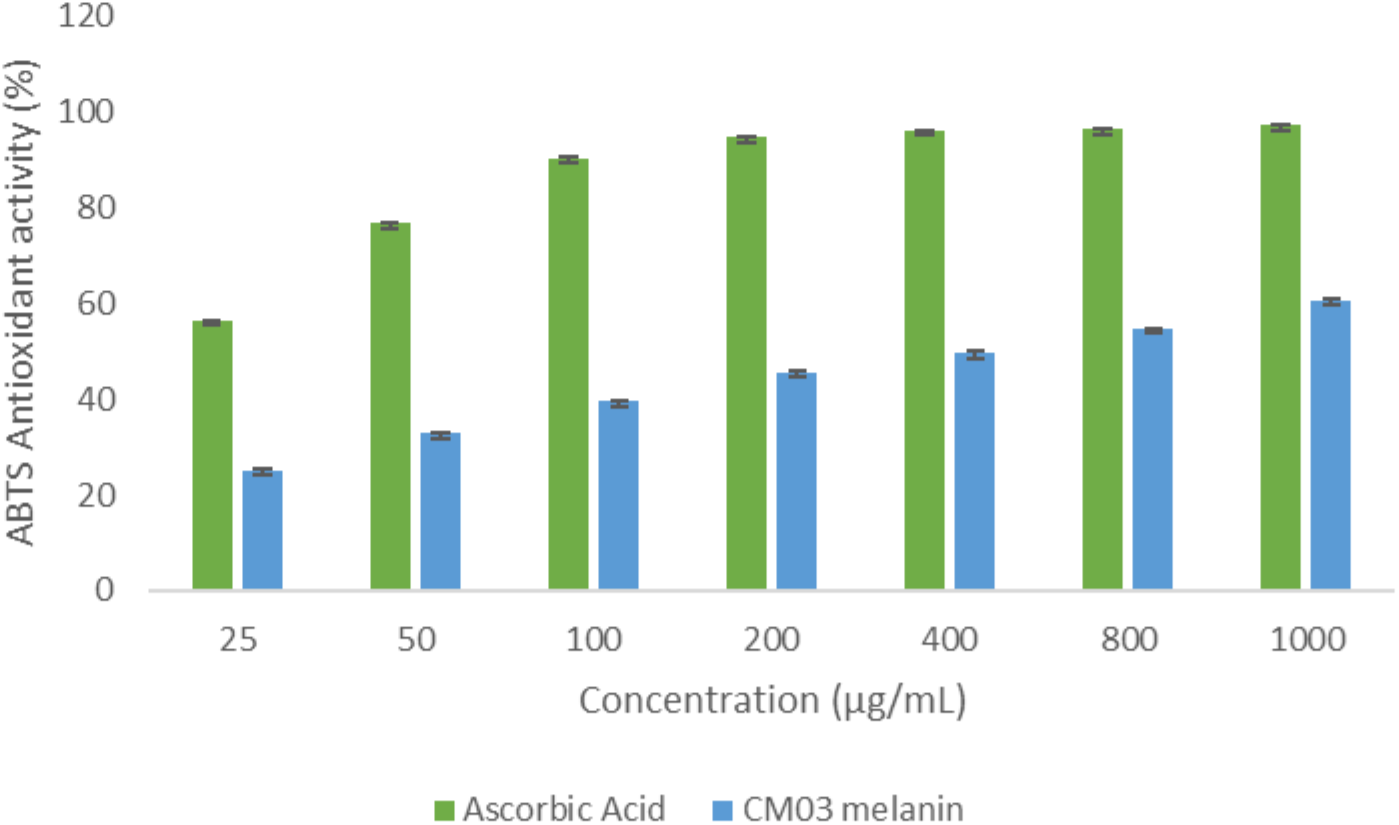
ABTS antioxidant activity of CM03 melanin.

### Ferric Reducing Antioxidant Power (FRAP) Assay

CMO3 melanin showed a clear, concentration-dependent increase in ferric-reducing antioxidant power (Figure 11). Fe²⁺-equivalent antioxidant activity rose from 0.23 ± 0.07 ng/mL at 6.25 µg/mL to 13.06 ± 0.16 ng/mL at 100 µg/mL, a roughly 57-fold increase across the tested range.

**Figure 11.**
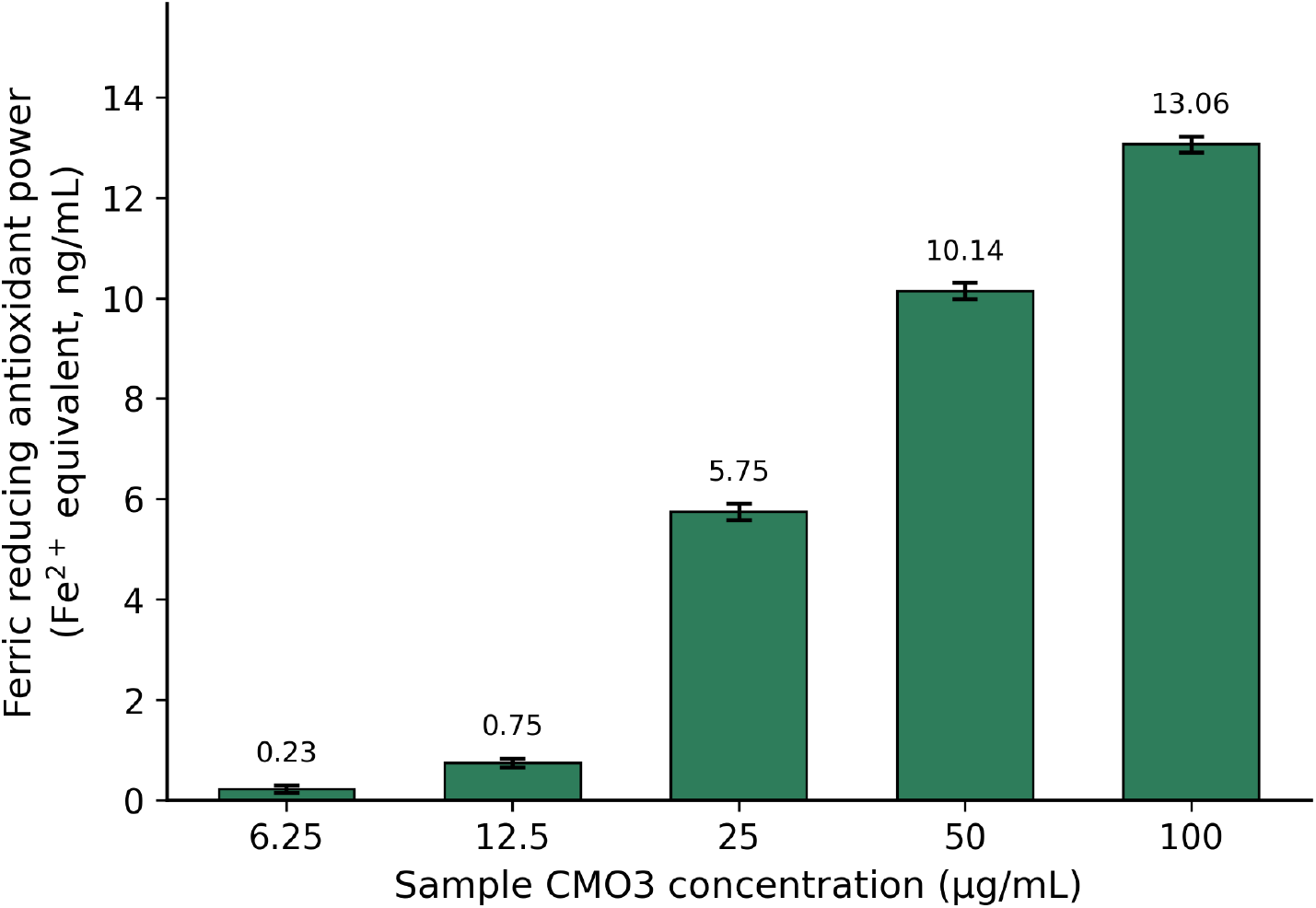
Ferric reducing antioxidant power (FRAP) of sample CMO3 across the tested concentration range, expressed as Fe²⁺ equivalents.

**Figure 12.**
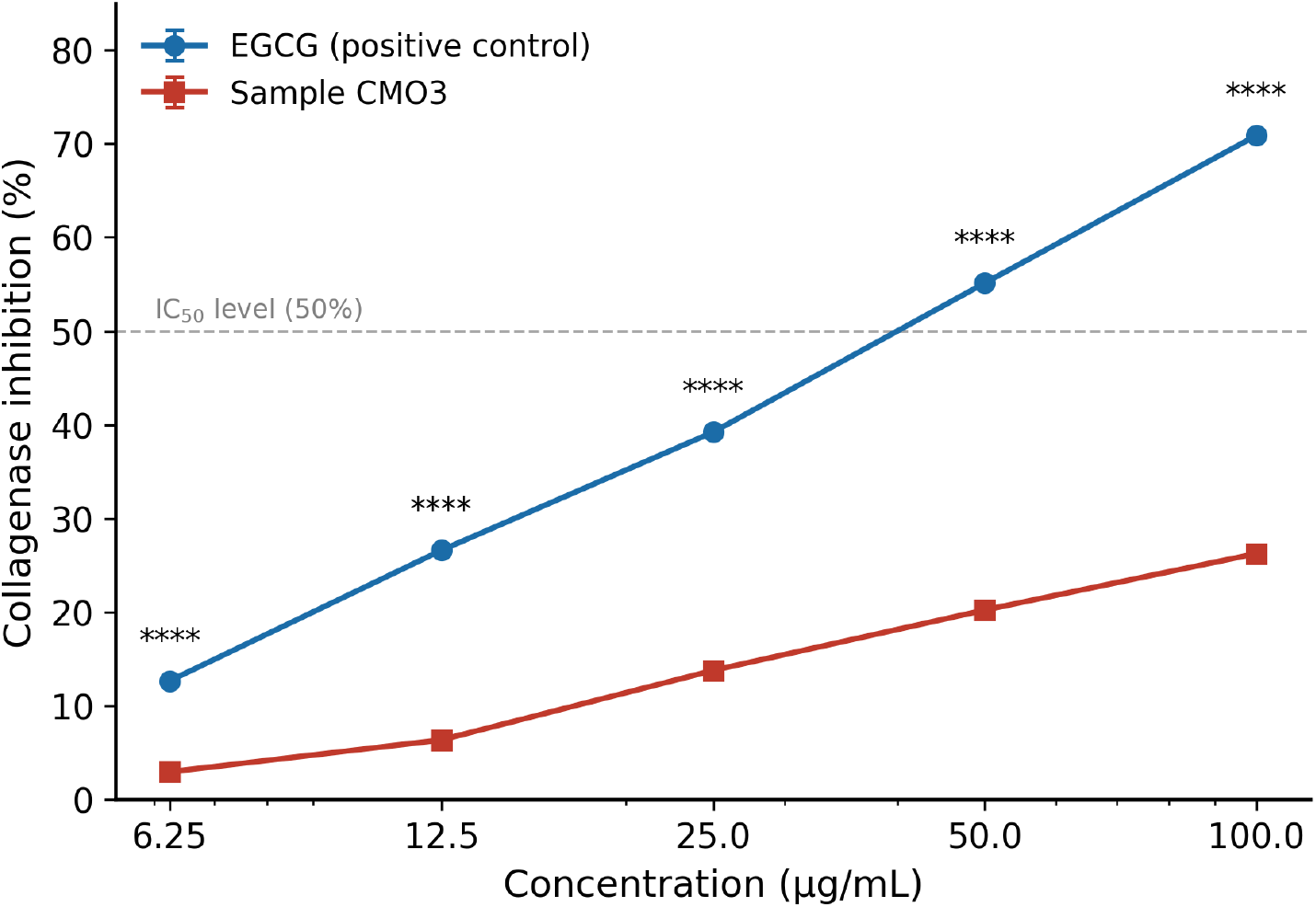
Dose-dependent collagenase inhibitory activity of sample CMO3 compared with EGCG (positive control). Data are mean ± SD (n = 3). **** p < 0.0001, unpaired t-test with Welch’s correction (CMO3 vs. EGCG at each concentration).

**Figure 13.**
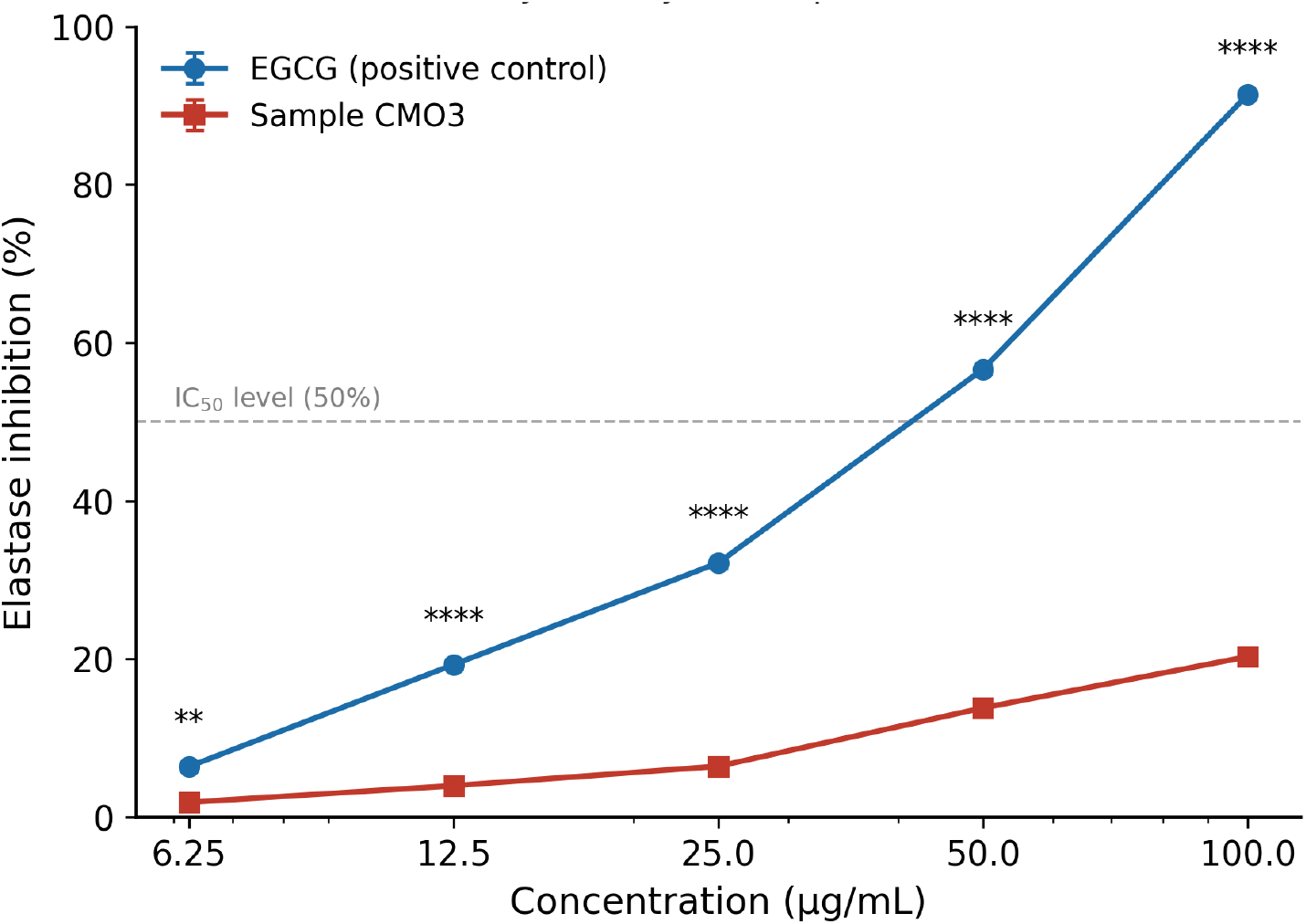
Dose-dependent elastase inhibitory activity of sample CMO3 compared with EGCG (positive control). Data are mean ± SD (n = 3). ** p < 0.01, **** p < 0.0001, unpaired t-test with Welch’s correction (CMO3 vs. EGCG at each concentration).

The FRAP assay provides a direct measure of the electron-donating (reducing) capacity of a sample, which is one of the principal mechanisms by which antioxidants neutralize reactive oxygen species and protect biomolecules — including skin collagen and elastin — from oxidative damage (Nishaa et al., 2012). CMO3 melanin exhibited a clear, concentration-dependent increase in ferric-reducing capacity across the tested range (6.25–100 µg/mL). This monotonic dose–response, together with the low variability among replicates (SD ≤ 0.16 ng/mL at all points), indicates that CMO3 contains electron-donating constituents with genuine, reproducible antioxidant activity under the conditions tested.

### Collagenase Enzyme Inhibitory (Anti-Ageing) Activity

CMO3 melanin had shown concentration-dependent increase in collagenase inhibition across the tested range of 6.25–100 µg/mL (Figure 12). Sample CMO3 showed markedly weaker inhibitory activity, increasing from 2.95 ± 0.07% at 6.25 µg/mL to only 26.25 ± 0.07% at 100 µg/mL compared to the control EGCG. One-way ANOVA confirmed a highly significant concentration-dependent effect for both EGCG (p < 0.0001) and CMO3 (p < 0.0001). At every concentration tested, inhibition by EGCG was significantly greater than that produced by CMO3 (unpaired t-test, p < 0.0001 at all five concentrations) indicating that CMO3 possesses substantially lower collagenase-inhibitory potency than the reference standard under the conditions tested.

Matrix metalloproteinases such as collagenase degrade the extracellular matrix components that maintain skin structure and elasticity, and their inhibition is a widely used in vitro correlate of anti-photoageing and anti-wrinkle potential (Park et al., 2010; Shanura Fernando et al., 2018). In this assay, CMO3 melanin exhibited only modest collagenase-inhibitory activity, reaching a maximum of 26.25% inhibition at 100 µg/mL. Nevertheless, the clear concentration-dependent trend observed for CMO3 (one-way ANOVA, p < 0.0001) suggests that the sample does possess intrinsic, if weak, anti-collagenase activity, and that higher concentrations than those tested here might reveal greater inhibition and permit IC₅₀ determination. Further work extending the concentration range beyond 100 µg/mL, would help to more completely characterize the cosmeceutical potential of this sample.

### Elastase Enzyme Inhibitory (Anti-Ageing) Activity

The CMO3 melanin produced a concentration-dependent increase in elastase inhibition across the tested range of 6.25–100 µg/mL (Figure 13). CMO3 showed markedly weaker inhibitory activity, increasing from 1.89 ± 0.16% at 6.25 µg/mL to only 20.31 ± 0.43% at 100 µg/mL compared to the EGCG standard. One-way ANOVA confirmed a highly significant concentration-dependent effect for both EGCG (p < 0.0001) and CMO3 (p < 0.0001). At every concentration tested, inhibition by EGCG was significantly greater than that produced by CMO3 (unpaired t-test with Welch’s correction, p < 0.01 at 6.25 µg/mL and p < 0.0001 at 12.5– 100 µg/mL), indicating that CMO3 possesses substantially lower elastase-inhibitory potency than the reference standard under the conditions tested.

Elastase is a serine proteinase that degrades elastin, a key extracellular matrix protein responsible for skin elasticity and resilience; inhibition of elastase activity is therefore widely used as an in vitro correlate of anti-ageing and anti-wrinkle potential (Senior et al., 1982; Sallenave et al., 1998). In this assay, CMO3 melanin exhibited only weak elastase-inhibitory activity, reaching a maximum of 20.31% inhibition at 100 µg/mL — its highest tested concentration. Testing the melanin sample at a higher concentration above 100 µg/mL might reveal greater inhibition. Taken together with the collagenase and elastase activity report, the highest tested concentration of melanin has shown only modest anti-ageing properties. Further studies using higher concentration of melanin needed for further confirmation.

### Cytotoxicity of melanin

CM03 nano melanin was found to show less cytotoxicity against the L929 mouse fibroblast cell line. Even at the higher concentration tested, the cell viability was found to be 96.10±0.43 % (Figure 14). Melanin did not change the morphology of the cells at the higher concentration tested. Results indicate the pigment’s noncytotoxic nature against normal cells. This could be advantageous for CM03 melanin nanoparticles to be used in cosmetic applications. Vijayan et al. (2017) reported that up to 200 ppm of bacterial melanin was found to be non-cytotoxic. Research on CM03 nano melanin reflects a similar, less cytotoxic nature of the pigment.

**Figure 14:**
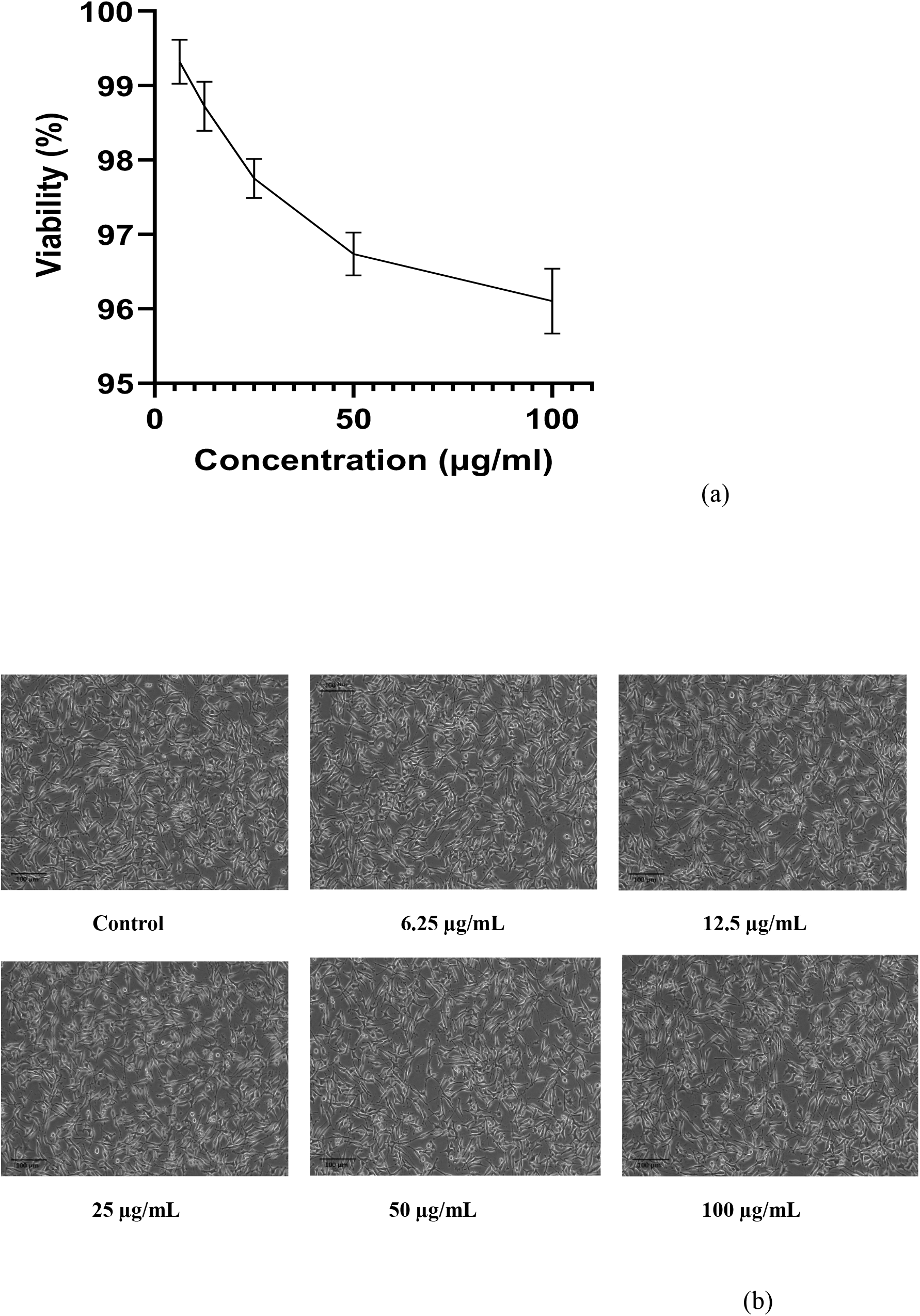
Cytotoxicity CM03 melanin nanoparticles. (a) concentration-dependent toxicity evaluation of melanin nanoparticles. (b) Effect of different concentrations of melanin on mouse fibroblast cell morphology.

## References

1. Bauer, A. W., Kirby, W. M. M., Sherris, J. C., & Turck, M. (1966). Antibiotic susceptibility testing by a standardized single disk method. American journal of clinical pathology, 45(4_ts), 493–496.

2. Bayram, S., Aygün, B., Karadayi, M., Alaylar, B., Güllüce, M., & Karabulut, A. (2023). Determination of toxicity and radioprotective properties of bacterial and fungal eumelanin pigments. International Journal of Radiation Biology, 99(11), 1785–1793.

3. Choi, K. Y. (2021). Bioprocess of microbial melanin production and isolation. Frontiers in bioengineering and biotechnology, 9, 765110.

4. Dalfard, A. B., Khajeh, K., Soudi, M. R., Naderi-Manesh, H., Ranjbar, B., & Sajedi, R. H. (2006). Isolation and biochemical characterization of laccase and tyrosinase activities in a novel melanogenic soil bacterium. Enzyme and microbial technology, 39(7), 1409–1416.

5. Dharmik, P. G., & Gomashe, A. V. (2013). Isolation, identification and antioxidant activity of melanin pigment from actinomycete (Streptomyces Species) Isolated from Garden Soil, Nagpur District, India. Int J Pure Appl Sci Technol, 18(1), 69–72.

6. Drewnowska, J. M., Zambrzycka, M., Kalska-Szostko, B., Fiedoruk, K., & Swiecicka, I. (2015). Melanin-like pigment synthesis by soil Bacillus weihenstephanensis isolates from Northeastern Poland. PloS one, 10(4), e0125428.

7. Ebrahimzadeh, M. A., Enayatifard, R., Khalili, M., Ghaffarloo, M., Saeedi, M., & Charati, J. Y. (2014). Correlation between sun protection factor and antioxidant activity, phenol and flavonoid contents of some medicinal plants. Iranian journal of pharmaceutical research: IJPR, 13(3), 1041.

8. El-Naggar, N. E. A., & El-Ewasy, S. M. (2017). Bioproduction, characterization, anticancer and antioxidant activities of extracellular melanin pigment produced by newly isolated microbial cell factories Streptomyces glaucescens NEAE-H. Scientific reports, 7(1), 1–19.

9. El-Naggar, N. E. A., & Saber, W. I. (2022). Natural melanin: current trends, and future approaches, with especial reference to microbial source. Polymers. 2022; 14: 1339.

10. Elsayis, A., Hassan, S. W., Ghanem, K. M., & Khairy, H. (2022). Optimization of melanin pigment production from the halotolerant black yeast Hortaea werneckii AS1 isolated from solar salter in Alexandria. BMC microbiology, 22(1), 92.

11. Ferraz, A. R., Pacheco, R., Vaz, P. D., Pintado, C. S., Ascensão, L., & Serralheiro, M. L. (2021). Melanin: Production from cheese bacteria, chemical characterization, and biological activities. International journal of environmental research and public health, 18(20), 10562.

12. Gonçalves, R. C. R., Lisboa, H. C. F., & Pombeiro-Sponchiado, S. R. (2012). Characterization of melanin pigment produced by Aspergillus nidulans. World Journal of Microbiology and Biotechnology, 28, 1467–1474.

13. Grigary, S., Umesh, M., & Mani, V. M. (2024). Isolation and characterization of polyhydroxyalkanoate producing halotolerant Bacillus subtilis SG1 using marine water samples collected from Calicut coast, Kerala. Journal of Applied Biology and Biotechnology, 12(2), 282–288.

14. Guo, L., Li, W., Gu, Z., Wang, L., Guo, L., Ma, S., … Chang, J. (2023). Recent advances and progress on melanin: From source to application. International journal of molecular sciences, 24(5), 4360.

15. Ito, S., Kolbe, L., Weets, G., & Wakamatsu, K. (2019). Visible light accelerates the ultraviolet A-induced degradation of eumelanin and pheomelanin. Pigment Cell & Melanoma Research, 32(3), 441–447.

16. Jalili, S., Pandey, R. P., & Kurian, N. K. (2022). Recent Insights in Melanin Research: From Extraction to Immense Applications of The Pigment. Journal of Survey in Fisheries Sciences, 140–146.

17. Jiang, H., Liu, N. N., Liu, G. L., Chi, Z., Wang, J. M., Zhang, L. L., & Chi, Z. M. (2016). Melanin production by a yeast strain XJ5-1 of Aureobasidium melanogenum isolated from the Taklimakan desert and its role in the yeast survival in stress environments. Extremophiles, 20, 567–577.

18. Jigna, C., & Gordhanbhai, P. M. (2022). Bacterial melanin with immense cosmetic potential produced by marine bacteria Bacillus pumilus MIN3.

19. Kamarudheen, N., Naushad, T., & Rao, K. V. B. (2019). Biosynthesis, characterization and antagonistic applications of extracellular melanin pigment from marine Nocardiopsis Sps. Indian J Pharm Educ Res, 53(2), 112–120.

20. Kiran, G. S., Jackson, S. A., Priyadharsini, S., Dobson, A. D., & Selvin, J. (2017). Synthesis of Nm-PHB (nanomelanin-polyhydroxy butyrate) nanocomposite film and its protective effect against biofilm-forming multi drug resistant Staphylococcus aureus. Scientific reports, 7(1), 9167.

21. Krumperman, P. H. (1983). Multiple antibiotic resistance indexing of Escherichia coli to identify high-risk sources of fecal contamination of foods. Applied and environmental microbiology, 46(1), 165–170.

22. Kurian, N. K., & Bhat, S. G. (2018). Data on the characterization of non-cytotoxic pyomelanin produced by marine Pseudomonas stutzeri BTCZ10 with cosmetological importance. Data in brief, 18, 1889–1894.

23. Mac Faddin, J. F. (1976). Biochemical Tests for Identification of Medical Bacteria 527.

24. Madhusudhan, D. N., Mazhari, B. B. Z., Dastager, S. G., & Agsar, D. (2014). Production and cytotoxicity of extracellular insoluble and droplets of soluble melanin by Streptomyces lusitanus DMZ-3. BioMed Research International, 2014.

25. Manivasagan, P., Venkatesan, J., Sivakumar, K., & Kim, S. K. (2013). Actinobacterial melanins: current status and perspective for the future. World Journal of Microbiology and Biotechnology, 29, 1737–1750.

26. Mbonyiryivuze, A., Nuru, Z. Y., Ngom, B. D., Mwakikunga, B. W., Dhlamini, S. M., Park, E., & Maaza, M. (2015). Morphological and chemical composition characterization of commercial sepia melanin.

27. Meredith, P., & Sarna, T. (2006). The physical and chemical properties of eumelanin. Pigment cell research, 19(6), 572–594.

28. Noman, E. A., Al-Gheethi, A., Al-Sahari, M., Mohamed, R. M. S. R., Crane, R., Ab Aziz, N. A., & Govarthanan, M. (2022). Challenges and opportunities in the application of bioinspired engineered nanomaterials for the recovery of metal ions from mining industry wastewater. Chemosphere, 308, 136165.

29. Nosanchuk, J. D., & Casadevall, A. (2003). The contribution of melanin to microbial pathogenesis. Cellular microbiology, 5(4), 203–223.

30. Nurhidayah, N., Diniatik, D., & Nurulita, N. A. (2022). Antioxidant Activity, Sun Protection Factor (SPF) and Total Phenolic and Flavonoid Contents from Purified Extract of Stelechocarpus buharol (BI.) Hook F. & Th. Leaves and its Classification with Chemometrics. International Journal of Nanoscience and Nanotechnology, 18(3), 179–185.

31. Pavan, M. E., López, N. I., & Pettinari, M. J. (2020). Melanin biosynthesis in bacteria, regulation and production perspectives. Applied microbiology and biotechnology, 104(4), 1357–1370.

32. Raman, N. M., & Ramasamy, S. (2017). Genetic validation and spectroscopic detailing of DHN-melanin extracted from an environmental fungus. Biochemistry and biophysics reports, 12, 98–107.

33. Rudrappa, M., Kumar, S., Kumar, R. S., Almansour, A. I., Perumal, K., & Nayaka, S. (2022). Bioproduction, purification and physicochemical characterization of melanin from Streptomyces sp. strain MR28. Microbiological Research, 263, 127130.

34. Saitou, N., & Nei, M. (1987). The neighbor-joining method: a new method for reconstructing phylogenetic trees. Molecular biology and evolution, 4(4), 406–425.

35. Sajjan, S. S., Anjaneya, O., Kulkarni, G. B., Nayak, A. S., Mashetty, S. B., & Karegoudar, T. B. (2013). Properties and functions of melanin pigment from Klebsiella sp. GSK. Korean J Microbiol Biotechnol, 41(1), 60–69.

36. Sangwan, A., & Jornet, J. M. (2022). Joint Communication and Bio-Sensing With Plasmonic Nano-Systems to Prevent the Spread of Infectious Diseases in the Internet of Nano-Bio Things. IEEE Journal on Selected Areas in Communications, 40(11), 3271–3284.

37. Singh, S., Nimse, S. B., Mathew, D. E., Dhimmar, A., Sahastrabudhe, H., Gajjar, A., … & Shinde, P. B. (2021). Microbial melanin: Recent advances in biosynthesis, extraction, characterization, and applications. Biotechnology Advances, 53, 107773.

38. Sun, S., Zhang, X., Chen, W., Zhang, L., & Zhu, H. (2016). Production of natural edible melanin by Auricularia auricula and its physicochemical properties. Food Chemistry, 196, 486–492.

39. Swift, J. A. (2009). Speculations on the molecular structure of eumelanin. International journal of cosmetic science, 31(2), 143–150.

40. Tamura, K., Dudley, J., Nei, M., & Kumar, S. (2007). MEGA4: molecular evolutionary genetics analysis (MEGA) software version 4.0. Molecular biology and evolution, 24(8), 1596–1599.

41. Tarangini, K., & Mishra, S. (2013). Production, characterization and analysis of melanin from isolated marine Pseudomonas sp. using vegetable waste. Res J Eng Sci, 2278, 9472.

42. Tarangini, K., & Mishra, S. (2014). Production of melanin by soil microbial isolate on fruit waste extract: two step optimizations of key parameters. Biotechnology Reports, 4, 139–146.

43. Thompson, C. R., Gerstman, B. S., Jacques, S. L., & Rogers, M. E. (1996). Melanin granule model for laser-induced thermal damage in the retina. Bulletin of mathematical biology, 58(3), 513–553.

44. Tran-Ly, A. N., Reyes, C., Schwarze, F. W., & Ribera, J. (2020). Microbial production of melanin and its various applications. World Journal of Microbiology and Biotechnology, 36, 1–9.

45. Vasanthabharathi, V., Lakshminarayanan, R., & Jayalakshmi, S. (2011). Melanin production from marine Streptomyces. African journal of biotechnology, 10(54), 11224.

46. Vijayababu, P., & Kurian, N. K. (2021). Melanin and its precursors as effective antiviral compounds: With a special focus on SARS CoV2. Mol Biol, 10, 1–3.

47. Vijayan, D., Young, A., Teng, M. W., & Smyth, M. J. (2017). Targeting immunosuppressive adenosine in cancer. Nature Reviews Cancer, 17(12), 709–724.

48. Wibowo, J. T., Kellermann, M. Y., Petersen, L. E., Alfiansah, Y. R., Lattyak, C., & Schupp, P. J. (2022). Characterization of an insoluble and soluble form of melanin produced by Streptomyces cavourensis SV 21, a sea cucumber associated bacterium. Marine Drugs, 20(1), 54.

49. Xu, C., Li, J., Yang, L., Shi, F., Yang, L., & Ye, M. (2017). Antibacterial activity and a membrane damage mechanism of Lachnum YM30 melanin against Vibrio parahaemolyticus and Staphylococcus aureus. Food Control, 73, 1445–1451.

50. Yabuuchi, E., & Ohyama, A. (1972). Characterization of “pyomelanin”-producing strains of Pseudomonas aeruginosa. International Journal of Systematic and Evolutionary Microbiology, 22(2), 53–64.

51. Zecca, L., Tampellini, D., Gatti, A., Crippa, R., Eisner, M., Sulzer, D., … & Gallorini, M. (2002). The neuromelanin of human substantia nigra and its interaction with metals. Journal of neural transmission, 109, 663–672.

52. Shanura Fernando, I. P., Asanka Sanjeewa, K. K., Samarakoon, K. W., Kim, H. S., Gunasekara, U. K. D. S. S., Park, Y. J., … & Jeon, Y. J. (2018). The potential of fucoidans from Chnoospora minima and Sargassum polycystum in cosmetics: Antioxidant, anti-inflammatory, skin-whitening, and antiwrinkle activities. Journal of Applied Phycology, 30(6), 3223–3232.

53. Park, K. J., Park, S. H., & Kim, J. K. (2010). Anti-wrinkle activity of Acanthopanax senticosus extract in ultraviolet B (UVB)-induced photoaging.Journal of Korean Society of Food Science and Nutrition, 39(1),42–46.

54. Sallenave, J. M., Xing, Z., Simpson, A. J., Graham, F. L., & Gauldie, J. (1998). Adenovirus-mediated expression of an elastase-specific inhibitor (elafin): a comparison of different promoters. Gene Therapy, 5(3), 352–360.

55. Senior, R. M., Campbell, E. J., Landis, J. A., Cox, F. R., Kuhn, C., & Koren, H. S. (1982). Elastase of U-937 monocytelike cells: comparisons with elastases derived from human monocytes and neutrophils and murine macrophagelike cells. The Journal of Clinical Investigation, 69(2), 384–393.

56. Nishaa, S., Vishnupriya, M., Sasikumar, J. M., Christabel, P. H., & Gopalakrishnan, V. K. (2012). Antioxidant activity of ethanolic extract of Maranta arundinacea L. tuberous rhizomes. Asian Journal of Pharmaceutical and Clinical Research, 5(4), 85–88.

